# A Sulfotransferase from a Gut Microbe Acts on Diverse Phenolic Sulfate Compounds, Including Acetaminophen Sulfate

**DOI:** 10.1101/2025.07.07.663557

**Authors:** Rylee Close, Schuyler Kremer, Madison Mitchem, Andrew Bellinghiere, Khemlal Nirmalkar, Chad R. Borges, Rosa Krajmalnik-Brown, Dhara D. Shah

## Abstract

Sulfonation is one of the two main phase II detoxification pathways in eukaryotes that transforms non-polar compounds into hydrophilic metabolites. Sulfotransferases catalyze these reactions by transferring a sulfo group from a donor to an acceptor molecule. Human cytosolic sulfotransferases use only 3’-phosphoadenosine 5’-phosphosulfate (PAPS) as a donor to sulfonate a variety of chemicals. Less understood are microbial aryl-sulfate sulfotransferases (ASSTs), which catalyze sulfo transfer reactions, without utilizing PAPS as a donor. Currently, the identity of physiological sulfo donor substrates remains unknown and sulfo acceptor substrates are underexplored. With this study, we aim to understand the potential contribution of a gut microbial enzyme to sulfonation chemistry by uncovering substrate preferences. Here, we show that a sulfotransferase (BvASST) from the prevalent gut microbe *Bacteroides vulgatus (now Phocaeicola vulgatus)* is a versatile catalyst that utilizes a wide range of phenolic molecules as substrates that are commonly encountered by the host. With this action, it has the ability to modulate concentrations of donor phenolic sulfates like acetaminophen sulfate, dopamine sulfate, p-coumaric acid sulfate, indoxyl sulfate, and p-cresol sulfate *in vitro*. Moreover, we report a large adaptability in the acceptor preferences with the evidence of sulfonation for many biologically relevant phenolic molecules including p-coumaric acid, p-cresol, dopamine, acetaminophen, tyramine, and 4-ethylphenol. These results suggest that such gut microbial enzymes may impact the detoxification of a variety of phenolic molecules in the host, which were previously thought to be solely detoxified via human sulfotransferases. However, further *in vivo* studies are necessary to understand potential contributions of ASSTs in host detoxification processes.

## INTRODUCTION

Humans have the ability to transform many endogenous and exogenous molecules via conjugation reactions which are considered Phase II detoxification pathways[1]. This process enhances the solubility of molecules, facilitating their metabolism and excretion. Sulfonation is one such major and prevalent conjugation chemistry. In humans, this chemistry is catalyzed by a multitude of cytosolic sulfotransferases, collectively known as hSULTs[2]. Sulfotransferases catalyze the transfer of a sulfo group from a donor molecule to an acceptor molecule. All known hSULTS studied so far show a stringent specificity towards the general sulfo group donor, 3’-phosphoadenosine 5’-phosphosulfate (PAPS). However, there is a divergence in hSULTs based on their acceptor preferences. These enzymes can sulfonate a variety of steroidal[3–5] and phenolic[6–8] metabolites that are relevant to human health. Additionally, A PAPS-dependent sulfotransferase, capable of converting the dietary steroidal metabolite cholesterol into cholesterol sulfate and regulating the concentrations of these molecules in the host, has recently been discovered in the commonly occurring gut microbe *Bacteroides thetaiotaomicron*[9, 10]. Products of such gut microbial metabolism can be absorbed by the host, undergo systemic circulation, or interact with epithelial cells[5]. Therefore, sulfonation of small molecules by gut microbial enzymes in the human gut might affect the rate of detoxification and pharmacokinetics of various molecules. These are some of the initial lines of evidence that potentially challenge the idea that sulfonation chemistry is exclusively catalyzed by hSULTS.

There is another known class of microbial sulfotransferases, called aryl-sulfate sulfotransferases (ASSTs), which are PAPS-independent and are known to sulfonate a variety of phenolic molecules[11–18] (Figure 1). We show that genes encoding ASSTs are present in the human gut microbiome, by analyzing genomes of the commonly occurring gut microbes and with datasets collected from human fecal samples from our previous study[19]. The variety of natural and synthetic acceptor molecules that interact with this class of enzymes has been previously investigated yet the identity of sulfo group donors for ASSTs still remain elusive. Unlike hSULTs, ASSTs are able to use a few different donor molecules, generally synthetic phenolic sulfates[11, 20–22]. Here, we identified previously unknown donors for an ASST from the prominent human gut microbe, *Bacteroides vulgatus (now Phocaeicola vulgatus)* and delved into donor and acceptor specificity of BvASST. Our results suggest that BvASST is able to modulate concentrations of a multitude of compounds of dietary, environmental, and pharmaceutical origin, as well as endogenous metabolites *in vitro*. Some of these molecules with health-related implications include, acetaminophen, tyramine, dopamine, p-cresol (4-methylphenol, pC), 4-ethylphenol (4-EP) and their sulfated forms.

**Fig. 1:**
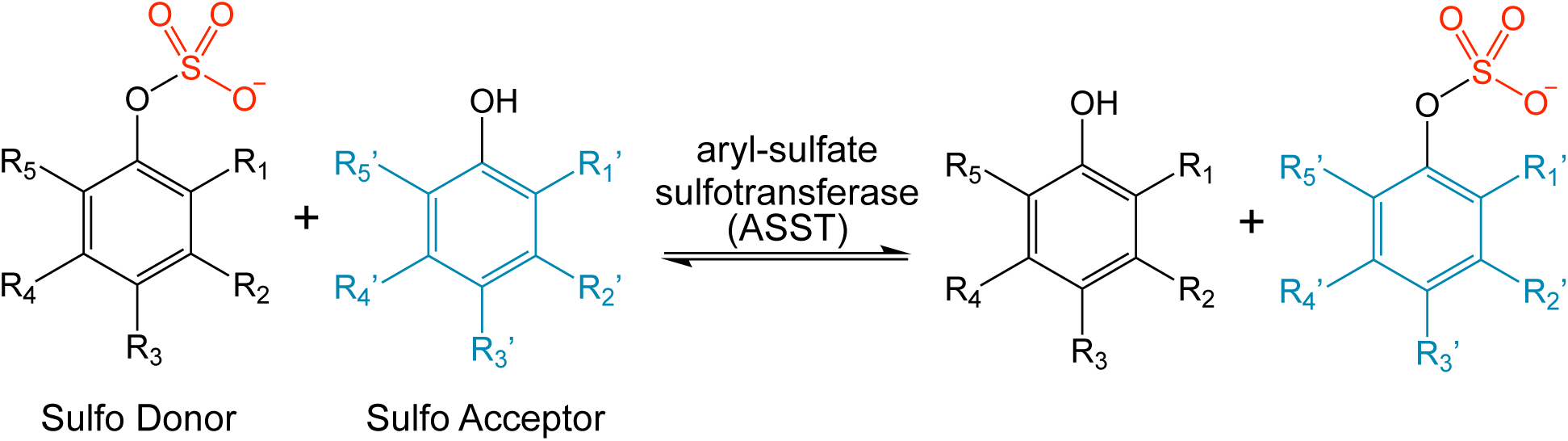
Schematic of the general reaction catalyzed by aryl-sulfate sulfotransferases (ASSTs).

In addition to substrate identification, we detected a sulfonated intermediate form of BvASST in the presence of our newly identified phenolic sulfo donors and determined the site of sulfonation to be His471 consistent with previously studied *E. coli* ASST[13]. We also showed that the stable sulfonated form of BvASST transfers sulfo group to various phenolic acceptor molecules and reverts to its original unsulfonated form that is ready for another round of catalysis. With this, we captured the entire catalytic cycle for BvASST in presence of newly identified donors and acceptors. The identities of substrates elucidated in this study provide the basis for understanding potential native reactions catalyzed by ASSTs in their natural ecosystem.

## MATERIALS AND METHODS

### Materials

The list of all chemical and materials utilized in this study is provided in Supplementary Table 1.

### Experimental methods

#### Mining annotated genes for aryl-sulfate sulfotransferases from genome databases and from human fecal sample metagenomics dataset

Genes annotated to encode aryl-sulfate sulfotransferases were found via Integrated Microbial Genomes and Microbiomes systems (IMG/M)[23, 24] and NCBI genome annotations[25] by searching with either EC 2.8.2.22 or with the name ‘aryl-sulfate sulfotransferase’ in commonly found 135 human gut microbes. To identify the presence or absence of aryl-sulfate sulfotransferase (*asst*) genes (KEGG Orthologs) in genomes of bacteria from human fecal samples, we used shotgun metagenomics data from our previous autism study [19] and looked for the *asst* genes in 38 children total (18 children with autism spectrum disorders (ASD) and 20 typically developing (TD) children). Additional details on the recruitment of participants, sample collection, sequence processing, and data analysis are available in a study by Nirmalkar et al.[19]. To identify *asst* genes from bacterial sequences, we used the marker gene-based tool MetaPhlAn4 (Metagenomic Phylogenetic Analysis, v3.1.0; https://github.com/biobakery/MetaPhlAn) and the taxonomic profiles of bacteria were identified with the ChocoPhlAn pangenome database[26]. Bacterial taxonomic profiles were used with HUMAnN3 (the Human Microbiome Project Unified Metabolic Analysis Network, v3.0; https://github.com/biobakery/biobakery/wiki/humann3) tool with UniRef90 and MetaCyc databases to calculate the bacterial gene abundance for metabolic pathways[27]. Further, gene family abundances from HUMAnN3 were converted to KEGG Orthologs (KOs) to analyze the differential relative abundance of KOs and *asst* gene.

#### Molecular cloning

*Bacteroides vulgatus* aryl-sulfate sulfotransferase (BvASST) was identified in preliminary gene product searches using Integrated Microbial Genomes & Microbiomes (IMG/M) database with EC:2.8.2.22, name= aryl-sulfate sulfotransferase. The gene for *Bacteroides vulgatus* aryl-sulfate sulfotransferase (Gene ID = 640760705, locus tag = BVU_RS02960, old locus tag = BVU_0583) was synthesized, and codon optimized for *E. coli* (Azenta Biosciences). The gene was cloned into pET-26b using restriction enzymes NcoI and SacI creating the construct pET-26b+BvASST (Supplementary Figure 2). A second construct was created without the signal peptide region (Supplementary Figures 4 & 5). For this, the *asst* gene from *B. vulgatus* was amplified with PCR using the forward primer 5’-CCATGGCCCGCGATGAAGAACAAGATT-3’ and the reverse primer 5’-GAGCTCTTACTGATTTTCCGGATA-3’. The resulting PCR product was cloned into pET-26b with restriction enzymes NcoI and SacI, creating the construct pET-26b+BvASSTnoSP. These constructs were then transformed into *E. coli* BL21(DE3) and *E. coli* NEB5α cells.

#### Protein expression and purification

*E. coli* BL21(DE3) cells harboring pET-26b+BvSTnoSP were grown in LB broth with 0.1 mM kanamycin at 37 °C until the OD_600_ reached 0.4, or 3.2 x 10^8^ bacterial cells/mL. At this point, the temperature was lowered to 20 °C and cultures were allowed to continue growing. Once an OD_600_ of 0.6, 4.8 x 10^8^bacterial cells/mL, was reached, cultures were induced with 0.1 mM IPTG and growth was continued at 20 °C overnight with shaking at 150 rpm. All the subsequent steps were carried out at 4 °C unless mentioned otherwise. Cells were harvested and resuspended in buffer containing 20 mM Tris, pH 8. One tablet of Roche cOmplete Mini EDTA-free protease inhibitor was added per liter of culture. Cells were lysed using sonication allowing for cycles of 30 seconds followed by 90 seconds of rest for 15 cycles at 60% duty cycle and tip output set at 6 with a Branson Sonifier 250 model 102C (CE). Sonicated cells were centrifuged at 18000*g* for 40 minutes. The collected supernatant then underwent ammonium sulfate precipitation. The protein of interest was precipitated out using 30% w/v ammonium sulfate which was then redissolved in 20 mM Tris buffer, pH 8 and filtered through a 0.2 µM PES membrane filter. This filtered solution was then diluted to 200 mL with 20 mM Tris buffer, pH 8 and loaded onto a pre-equilibrated Cytiva HiScreen Capto DEAE column (0.77 x 10 cm, 4.7 mL) at the flow rate of 1.5 mL/min. The column was then washed with 5 column volumes of buffer containing 20 mM Tris, 50 mM NaCl, pH 8 to remove contaminating proteins. BvASST was eluted with a linear salt gradient from 20 mM Tris, 50 mM NaCl, pH 8 to 20 mM Tris, 500 mM NaCl, pH 8 at the flow rate of 1 mL/min. The eluted protein was then buffer exchanged to 20 mM Tris, pH 8 and concentrated with 30 kDa molecular weight cut-off (MWCO) centrifugal filters from Cytiva. The protein was stored at – 80 °C. This purified protein was used for all enzymatic assays. For MS (Mass spectrometry) analysis, the protein was further concentrated and loaded onto a Cytiva HiPrep 16/60 Sephacyrl S-300 (120 mL) size exclusion column (SEC) and eluted with a buffer containing 20 mM Tris, 50 mM NaCl, pH 8 at a flow rate of 0.5 mL/min. Fractions with BvASST were pooled and concentrated with 30 kDa MWCO spin filters.

#### Enzyme activity assays

Enzyme activity assays contained 1 mM of sulfo donor, 1 mM of sulfo acceptor, and 4 µM of BvASST. All assays were conducted in in 20 mM Tris, pH 8. Absorbance based assays were performed individually in cuvettes with an Agilent Cary 3500 UV-Vis spectrophotometer and fluorescence-based assays were conducted in black 96 well plates with a Biotek Synergy HTX multi-mode plater reader in a kinetic mode. Absorbance based assays utilized p-nitrophenyl sulfate (pNPS) as a sulfo donor and were monitored at 405 nm for the production of p-nitrophenol (pNP) whereas fluorescence-based assays relied on 4-methylumbelliferyl sulfate (4-MUS) as a sulfo donor and were monitored at 360 excitation/460 emission for production of 4-methylumbelliferone (4-MU) using a standard curve. All assays were carried out at 37 °C and at a final volume of 100 µL for absorbance-based assays or 300 µL for fluorescence-based assays. Absorbance-based assays were monitored for 30 minutes whereas fluorescence-based assays were monitored for 200 minutes. All reactions were initiated with the addition of acceptor molecules, which allowed the incubation of enzyme with sulfo group donors. Initial velocities were calculated from the linear portions of the plots of product concentration over time.

#### Incubation of BvASST with various sulfo group donors to detect sulfonated BvASST

50 µM BvASST was incubated with either 0.1 or 1 mM of respective sulfo donor for 20 minutes at 37 °C in 20 mM Tris, pH 8. After incubation, these samples were spin filtered (30 kDa, MWCO spin filters) and transferred to a final buffer consisting of 5 mM ammonium acetate, pH 7.5 and immediately analyzed via intact mass spectrometry.

#### Incubation of sulfonated BvASST with various sulfo group acceptors to detect unsulfonated BvASST

50 µM BvASST was incubated with 50 µM pNPS at 37 °C for 20 minutes in 20 mM Tris, pH 8 to produce sulfonated BvASST. Sulfonated BvASST was then incubated with a variety of sulfo acceptors: p-coumaric acid (pCA), 4-ethylphenol (4-EP), dopamine, p-cresol, acetaminophen, tyramine, and a control without acceptor. Samples were incubated at 37 °C for 2 hours in 20 mM Tris, pH 8, before spin filtered (30 kDa, MWCO spin filters) and transferred to a final buffer of 5 mM ammonium acetate, pH 7.5. Control samples were processed similarly but not incubated with any substrate (without donor and acceptor). Samples were kept on ice and immediately analyzed via mass spectrometry. Intact mass analysis was carried out as detailed in the ‘*Intact protein mass spectra’* methods section.

#### Intact protein mass spectrometry

BvASST spectra were obtained on a Bruker Maxis 4G quadrupole-time of flight instrument with an ESI ion source connected to an Ultimate 3000 capillary HPLC equipped with an autosampler. Sample preparation included dilution with 0.1% (v/v) trifluoroacetic acid (TFA) to bring individual samples to one micromolar concentration, of which 2 µL was injected. Specimens were initially introduced via the loading pump at 10 µL/min with the solvent at an 80/20 mixture of 0.1% Formic acid and acetonitrile respectively for four minutes through an Optimize Technologies Protein Captrap (Cat number 10-04814-TM). After four minutes, a valve switch routed the capillary pump flow over the captrap, which carried the protein from the captrap into the mass spectrometer at 3 µL/min with a mixture of 0.1% Formic Acid (Solvent A) and 100% Acetonitrile (Solvent B) at a composition of 65%/35% (A/B), which was held until minute 4.6 when the stepwise gradient was changed to 55%/45% (A/B) over 0.1 min. At minute 6.6 the solvent composition was adjusted to 35%/65% (A/B) over 0.1 min, which then ran until 7.5 minutes when the composition was changed to 20%/80% (A/B) over 0.1 min. Finally, at 9.6 minutes the solvent composition was returned to the baseline of 65%/35% (A/B) and remained that way for the rest of the ten-minute run. The Bruker Maxis 4G ESI-Q-TOF was set in positive ion mode over a m/z range of 300-3000. The potential on the inlet capillary was set to 4000 V and end plate offset was 500 V. Nebulizer nitrogen was 3 bar and the dry gas nitrogen was set to 4.0 L/min at 225 °C. The sampler rate of the digitizer was 4GHz with a spectra rate summation of 1Hz. Data analysis was performed using Bruker Daltronic Compass Software wherein spectra were averaged from 4.6 to 7 minutes. Baseline subtraction of this summation was done at 0.7 flatness and then subsequently the raw mass spectra were charge deconvoluted. Following deconvolution another baseline subtraction was performed to visualize peaks for analysis.

#### Trypsin digestion of BvASST and mass spectrometry

The ASST specimens were reduced, alkylated, digested with trypsin then analyzed by liquid-chromatography coupled to either collision induced dissociation (CID) or electron transfer dissociation/higher-energy collisional dissociation (EThcD)-based tandem mass spectrometry (LC-MS/MS). Reduction, alkylation, and tryptic digestion was performed with a modified version of the S-Trap Protocol available on the manufacturer’s website as a micro spin column digestion protocol (Detailed in Supplemental Methods).

#### CID-Based LC-MS/MS

The trypsin-digested products of a 70 µg ASST sample were separated on an Ultimate 3000 liquid chromatography system equipped with a BIOshell™ A160 Peptide C18 column (150 mm X 0.2 mm, 2.7 µm particle diameter size (Sigma-Aldrich, 67206-u)) and eluted into a Bruker Maxis 4G quadrupole-time-of-flight mass spectrometer. Auto MS/MS was performed using a scheduled protocol list for 1263.5 to 1263.90 with CID energy set to scale between 30 and 50 eV, based on charge state. Details on the MS/MS protocol are included in the Supplemental Information section.

#### EThcD-Based LC-MS/MS

Peptides from trypsin-digested ASST were injected into a Orbitrap Fusion Lumos through an Ultimate 3000 using a C18 Easy Spray HPLC column (75 µm X 500 mm, 2 µm particle size (ThermoFisher, ES903)). Analysis by MS/MS was done using a target inclusion list for 1262.8698^3+^ and 1263.5381^3+^ with electron transfer dissociation set at 50 ms reagent time and a supplemental activation for HCD set at 20%. (Details in Supplemental Methods). Fragmentation carried out in this way tends to produce *c* and *z* ions in addition to *b* and *y* ions that are commonly produced by conventional collision-induced dissociation (CID)[28].

#### Mass spectrometry data processing

Theoretical fragment masses were calculated using Protein Prospector (https://prospector.ucsf.edu). Carbamidomethylation of cysteine was set as a fixed modification to account for alkylation by iodoacetamide. Additionally, sulfohistidine was set as a fixed modification on His471 using the User-Specified Elemental Composition feature to account for the 80 Da mass shift associated with sulfonation. Data from Orbitrap EThcD runs were analyzed using the FreeStyle Software from ThermoFisher Scientific. MS/MS scans were averaged over the chromatographic elution profile of m/z 1263.5670^3+^ to generate an information-rich MS/MS spectrum. Masses of observed monoisotopic peaks from fragment ions were compared against calculated fragment masses to calculate delta and ppm values. Assignments were made for observed fragments within ± 0.007 m/z units or ± 10 ppm from their theoretical (calculated) *m/z* values. Assignments are documented in Supplementary File 1, as well as in the MS/MS spectrum used to make those assignments.

#### HPLC analysis of BvASST catalyzed reactions

To detect production of sulfated acceptor molecules, we analyzed the samples using HPLC. All enzymatic reactions were analyzed with the Waters e2695 separation module using a Synergi 4 μm Hydro-RP 80 A 250 x 4.6 mm column, under gradient conditions using buffer A containing methanol with 0.1% formic acid and buffer B containing water with 0.1% formic acid. The optimized gradient method started with 5% A: 95% B for 0–5 min, to 20% A:80% B from 5–19 min, this changed to 80% A:20% B from 19–28 min. This combination was held from 28–35 min. After the hold for 7 min, the mixture was changed to 5% A:95% B from 35–37 min, which was then converted to starting condition of 5% A: 95% B from 37–50 min at a flow rate of 1 mL/min. All HPLC samples were monitored by a photo diode array detector. Specific wavelengths used for data analysis were 243 nm, 265 nm, 276 nm, and 315 nm. All enzymatic reactions and controls (n=3) were filtered through 10 kDa MWCO spin filters and stored at − 80 °C before being analyzed by HPLC. All products from enzyme catalysis were identified by comparing their retention times to authenticated standards purchased commercially.

#### Enzyme concentration variation experiments

Enzyme activity assays were conducted with various acceptors and pNPS as a donor. For each reaction using a unique acceptor and the donor pNPS, BvASST concentrations were varied from 0 µM to 8 µM in 2 µM increments. Concentrations of pNPS and acceptor were held at 1 mM. Reactions were carried out in 96 well plates in 20 mM Tris, pH 8 to a final volume of 300 µL for 150 min at 37 °C. All reactions were initiated with the addition of acceptor molecules. Initial velocities were measured as the rate of product formation over time, which is µM pNP formation per minute from the linear portions of the progress curves.

#### Steady-state kinetics

Steady state kinetic parameters were measured with activity assays mentioned above by varying either donor concentrations or acceptor concentrations.

##### a) Varying acceptor concentrations with a fixed concentration of pNPS as a donor

Rates of BvASST catalyzed reactions were measured with 1 mM pNPS as the sulfo group donor and different acceptor molecules at varying concentrations. These are p-coumaric acid (0.039 µM to 2.25 mM), dopamine (0.198 µM to 3.38 mM), acetaminophen (0.198 mM to 10 mM), 4-ethylphenol (0.250 mM to 5 mM), p-cresol (25 µM to 1.6 mM), and caffeic acid (12.5 µM to 500 µM). Different concentration ranges were used for each molecule based on the concentration needed to achieve the maximum velocity with BvASST. All reactions were carried out in 20 mM Tris, pH 8 with 4 µM of BvASST at 37 °C for 30 min and monitored by measuring the absorbance of the product pNP, at 405 nm spectrophotometrically. Reactions were started with the addition of acceptor molecule after allowing the donor and enzyme to equilibrate. Initial rates (V_o_) were measured by fitting the initial time points data to linear regression. These calculated initial rates were then plotted against the concentrations of each acceptor molecule separately. From the hyperbolic fit of these datasets, kinetic parameters were determined for each acceptor molecule. For volatile acceptors (p-cresol, 4-ethylphenol), reactions were catalyzed similarly but these were incubated in clear 96 well plates with covers.

##### b) Varying acceptor concentrations with a fixed concentration of 4-MUS as a donor

Rates of BvASST catalyzed reactions were measured with 1 mM 4-MUS as the sulfo donor and different acceptor molecules at varying concentrations, including p-coumaric acid (50 µM to 1.5 mM), dopamine (62.5 µM to 1 mM), acetaminophen (125 µM to 2 mM), 4-ethylphenol (125 µM -4 mM), p-cresol (125 µM – 4 mM), and caffeic acid (3.1 µM – 200 µM). Different concentration ranges were used for each molecule based on the concentration needed to saturate the enzyme velocity. All reactions were carried out in 20 mM Tris, pH 8 with 4 µM of BvASST at 37 °C for 200 min and monitored by measuring the fluorescence of the product, 4-MU (360 excitation/ 460 emission) in black 96 well plates. Initial rates (V_o_) were measured by fitting the initial time points data to linear regression. These calculated initial rates were then plotted against the concentrations of each acceptor molecule separately. From the hyperbolic fit of these datasets, kinetic parameters were determined for each acceptor molecule. Reactions were started with the addition of acceptor molecule after allowing the donor and enzyme to equilibrate. All steady state kinetics data were fitted with Michaelis–Menten equation (equation 1).

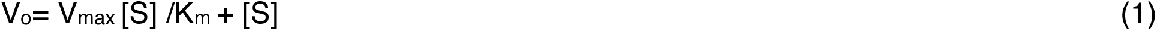

## RESULTS

### Putative genes annotated to encode microbial aryl-sulfate sulfotransferases are prevalent in gut microbes

To explore the possibility of sulfonation by microbial aryl-sulfate sulfotransferases being part of gut microbiota transformations, we investigated the genomes of commonly found gut microbes. As seen in Supplementary Figure 1, various gut bacterial strains, spanning multiple genera known to inhabit human GI tract, had genes annotated to encode aryl-sulfate sulfotransferases. Many displayed multiple putative *asst* genes within their genomes that were not direct copies. Particularly, microbes of the genus *Sutterella* displayed anywhere from 2 to 17 *asst* genes[29]. Separate from available genome databases, we identified putative genes annotated as aryl-sulfate sulfotransferases in a metagenome dataset from human fecal samples[19], where we found the presence of *asst* genes in 8 out of 38 samples (Supplementary Table 2). This analysis was not to compare abundances, but to inquire about the presence of these genes in fecal samples. Although these genes have not been highly studied and hence might be lacking in many databases, we were able to confirm their presence in about 20% of our previously sequenced samples[19].

### The sulfonation state of BvASST in the presence of various phenolic molecules, both sulfated and non-sulfated, reveals the identities of sulfo group donors and acceptors

Despite the high diversity of gut microbial *asst* genes, for this study, we chose to focus on an annotated *asst* gene from *Bacteroides vulgatus ATCC 8482 (now Phocaeicola vulgatus)* for several reasons: 1) the prevalence of *Bacteroides* in the human gut microbiome, 2) the presence of a unique lipoprotein signal peptide identified in the translated protein through *in silico* analysis, 3) existing studies on *Bacteroides thetaiotaomicron* that describe a different class of sulfotransferases[9, 10], and 4) the reduced complexity in selecting an organism with only one annotated *asst* gene, facilitating future mechanistic studies. Our initial attempts to purify aryl-sulfate sulfotransferase (BvASST) from *Bacteroides vulgatus* resulted in insoluble protein aggregates (Supplementary Figure 2b). Following this, we used SignalP 6.0[30] to determine if a signal peptide was present on BvASST. The program predicted that the enzyme contained a lipoprotein signal peptide (SEC/SPII) with a probability of 0.93 (Supplementary Figure 3a). With the removal of lipoprotein signal peptide and an addition of pelB leader sequence, BvASST was potentially untethered from the membrane and was purified in a soluble form (Supplementary Figure 4) when expressed in *E. coli*. This form of the enzyme was used in all subsequent experiments, particularly because the presence of the pelB sequence had little to no impact on BvASST activity (Supplementary Figures 5 and 6).

All previous studies characterizing ASSTs have been largely confined to utilizing synthetic molecules like p-nitrophenyl sulfate (pNPS) and 4-methylumbelliferyl sulfate (4-MUS) as sulfo donors[12, 16, 22], which are not found in physiological conditions. Other than these synthetic molecules, the identity of sulfo donors for ASSTs continues to remain elusive, whereas mammalian sulfotransferases are known for their stringent specificity to use PAPS as a sulfo group donor[2]. To identify potential sulfo donors with known physiological relevance to human health, we leveraged the catalytic mechanism that is characteristic of this class of enzymes known as ping-pong bi-bi[12, 16]. Within this mechanism, the two substrates, identified as the donor of the sulfo group and the acceptor of the sulfo group, interact with the enzyme in separate and successive stages rather than concurrently (Figure 2). Initially, the sulfo group donor attaches to the enzyme’s active site and transfers its sulfo group to the enzyme, departing in a desulfated form. This first half reaction temporarily converts the enzyme into a sulfonated intermediate (Figure 2). In the second half reaction, the sulfonated enzyme intermediate can now bind and transfer a sulfo group to an acceptor molecule to generate sulfated acceptor. During this process, the enzyme reverts to its original unsulfonated form that is ready to start another catalytic cycle.

**Fig. 2:**
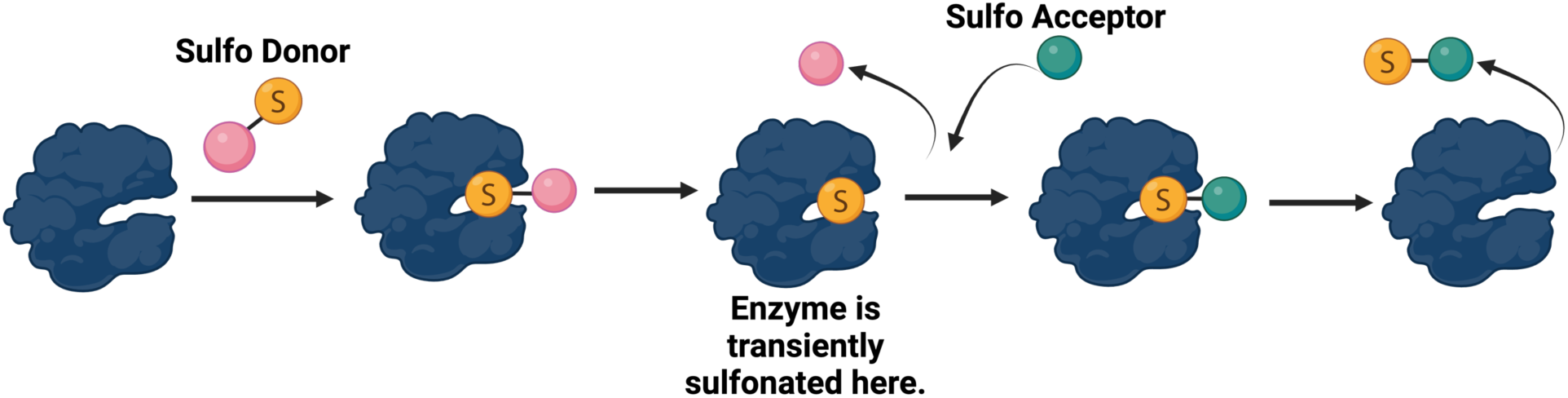
Predicted sulfonation state of BvASST throughout the course of the reaction[13, 31].

Taking advantage of this rection mechanism, we adapted an intact protein mass-spectrometry based assay from Malojcic et al.[13] and developed a screen to identify phenolic sulfo group donors and acceptors for BvASST. Initially, we measured the intact molecular mass of the protein with and without incubation with various sulfo group donor molecules. The unsulfonated form of the enzyme has a molecular mass of 68,267 Da, and when the enzyme adopts an intermediate sulfonated form a mass shift of +80 Da is expected. To screen sulfo group donors, BvASST was incubated with various phenolic sulfate molecules encountered by humans. Additionally, due to the predicted reversibility of ASST catalyzed reactions, a sulfated acceptor molecule produced by this reaction has the potential to act as a sulfo donor in the reverse direction. With this rationale, we tested commercially available sulfated forms of previously known acceptors known to interact with other microbial ASSTs[11, 14–18, 22, 32–34]. These molecules include p-coumaric acid sulfate (pCAS), acetaminophen sulfate (acetS), indoxyl sulfate (indoxylS), p-cresol sulfate (pCS), dopamine sulfate (dopamineS), serotonin sulfate (serotoninS), and PAPS. In addition, we incubated BvASST with synthetic sulfo donors like p-nitrophenyl sulfate (pNPS) and 4-methylumbelliferyl sulfate (4-MUS). Interestingly, in the presence of most of the above-mentioned phenolic sulfates, we found a mass shift of +80 Da to produce a peak representing the sulfonated intermediate of BvASST with an intact mass of 68,347 Da (Figure 3), demonstrating that sulfonation reactions were taking place. We did not detect a sulfonated form of BvASST when using serotonin sulfate and PAPS as sulfo donors (Figure 3), suggesting that neither molecule functions as a sulfo group donor for this enzyme. Although PAPS is a common sulfo donor for all human SULTs, it is not an aryl sulfate, which may explain its ineffectiveness with BvASST. Serotonin sulfate, despite being an aryl sulfate, also failed to transfer a sulfo group possibly due to its amine characteristic. To better understand the structural and chemical requirements of sulfo donors for BvASST, additional testing with molecules structurally similar to serotonin sulfate is needed. These results showed that exogenous and endogenous phenolic sulfates with biological relevance to humans are sulfo donors for BvASST. In agreement with other ASSTs, BvASST was also sulfonated in the presence of synthetic sulfo donors (Figure 3).

**Fig. 3:**
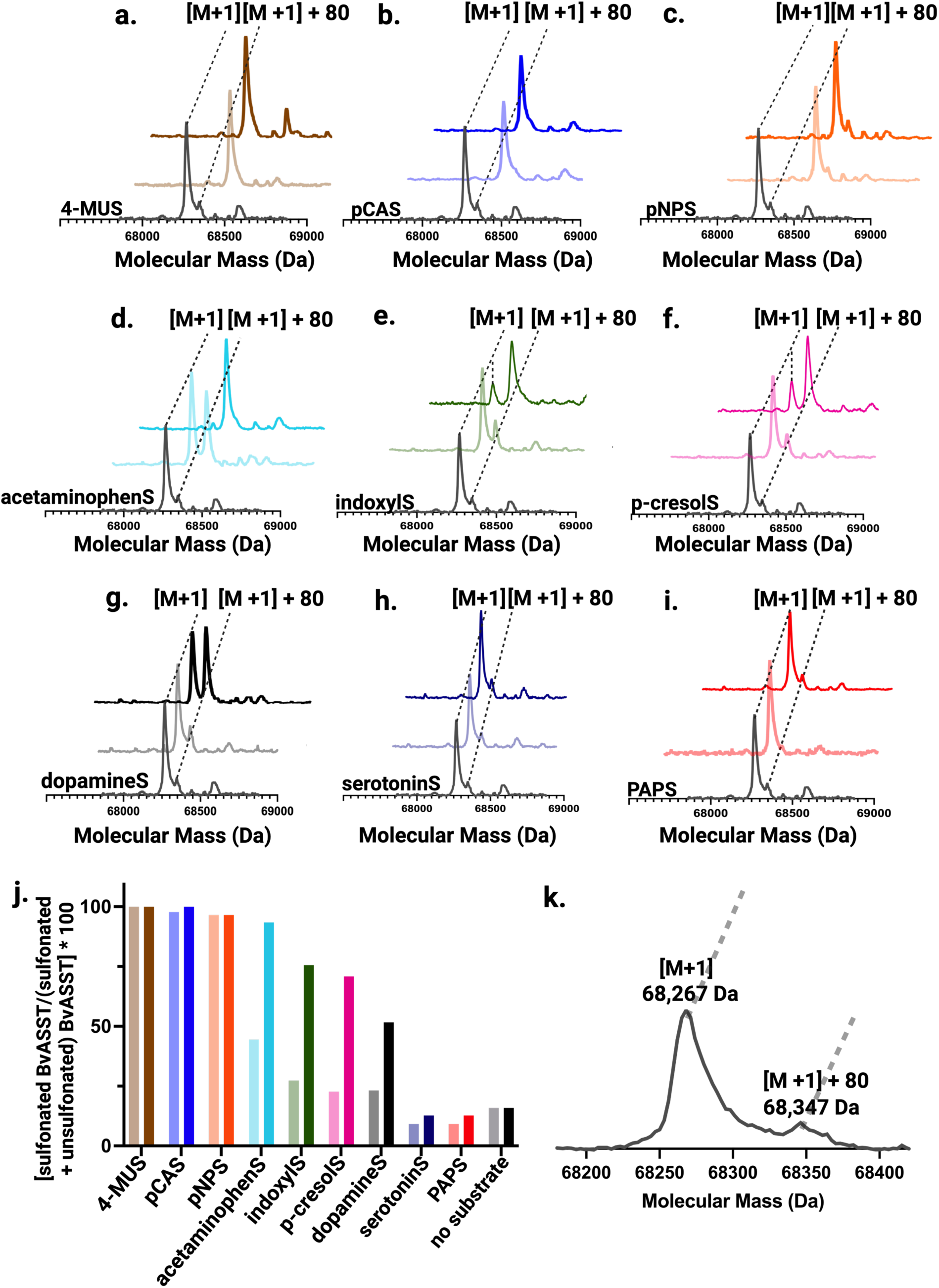
BvASST undergoes sulfonation by several phenolic sulfates. Intact protein mass-spectra of sulfonated (([M+1]+80)) and unsulfonated ([M+1]) forms of BvASST with various donors are shown above. In each panel, the black, baseline spectrum represents BvASST without the addition of the donor substrate, which contains a fraction of enzyme in a sulfonated form. The lighter color spectrum in each panel represents mass-spectrum of intact BvASST after the incubation with 100 µM of donor and the darker shade represents mass-spectrum of intact BvASST after the incubation with 1 mM of donor. All incubations were carried out at 37 °C for 20 minutes. Here, BvASST spectra are captured with, **a.** 4-methylumbelliferyl sulfate (4-MUS) **b.** p-coumaric acid sulfate (pCAS), **c.** p-nitrophenyl sulfate (pNPS), **d.** acetaminophen sulfate (acetS), **e.** indoxyl sulfate (indoxylS), **f.** p-cresol sulfate (pCS), **g.** dopamine sulfate (dopamineS), **h.** serotonin sulfate (serotoninS), **i.** PAPS, **j.** Schematic of qualitative trends in enzyme sulfonation across various donors and concentrations where colors and concentrations correspond to spectra shown in panels a-i, estimated as a percentage of ratio: ([M+1] + 80)/ ([M+1] + ([M+1] + 80]))*100 or [sulfonated BvASST/ (sulfonated + unsulfonated) BvASST]*100. Here, 4-MUS is 4-methylumbelliferyl sulfate, pCAS is p-coumaric acid sulfate, pNPS is p-nitrophenyl sulfate, acetaminophenS is acetaminophen sulfate, indoxylS is indoxyl sulfate, p-cresolS is p-cresol sulfate, dopamineS is dopamine sulfate, and serotoninS is serotonin sulfate. **k.** Zoomed in baseline spectrum of purified BvASST depicting two peaks without any substrate. The major peak with molecular mass of 68,267 Da represents unsulfonated ([M + 1]) form of BvASST whereas the minor peak (≈ 16%) with molecular mass of 68,347 Da represents sulfonated ([M+ 1] + 80]) form of BvASST.

Additionally, the screen to identify phenolic sulfo donors was carried out with two separate concentrations of donor molecules against BvASST, 100 µM and 1000 µM (1 mM) (Figure 3). 4-MUS (Figure 3a), pCAS (Figure 3b), and pNPS (Figure 3c) fully sulfonated BvASST at only 100 µM as apparent by the complete shift of the mass peak [M+1] (68,267 Da) to [M+1] + 80 (68,347 Da) (Figure 3a-3c, Figure 3k, spectra with lighter shades) with no further increase in [M+1] + 80 (68,347 Da) peak which represents the sulfonated BvASST when donor concentrations were increased to 1 mM (Figure 3a-3c, spectra with darker shades). In contrast, BvASST was only partially sulfonated at 100 µM as apparent with the presence of two peaks ([M+1] which is unsulfonated form and [M+1] + 80 which is sulfonated form) with acetaminophen sulfate (Figure 3d, lighter shade spectrum), indoxyl sulfate (Figure 3e, lighter shade spectrum), p-cresol sulfate (Figure 3f, lighter shade spectrum), and dopamine sulfate (Figure 3g, lighter shade spectrum). When incubated with 1 mM, there was an increase in the mass peak ([M+1] + 80, 68,347 Da) representing sulfonated BvASST with concomitant decrease in the mass peak ([M+1], 68,267 Da) corresponding to unsulfonated BvASST in the presence of acetaminophen sulfate (Figure 3d, darker spectrum), indoxyl sulfate (Figure 3e, darker spectrum), p-cresol sulfate (Figure 3f, darker spectrum), and dopamine sulfate (Figure 3g, darker spectrum). We did not observe any mass shift of [M+1] peak (unsulfonated form) where BvASST was incubated with either serotonin sulfate (Figure 3h) or PAPS (Figure 3i), indicating that these molecules could not transfer sulfo group to BvASST and hence can’t be donors for BvASST. These differences in sulfonation process via various molecules are likely driven by the donor preferences of BvASST.

As per predicted mechanism, the sulfonated enzyme intermediate (sulfonated BvASST) transfers sulfo group to potential acceptors and returns to its original form (unsulfonated BvASST) (Figure 2). To screen possible sulfo acceptors, BvASST was first converted to its sulfonated form by the addition of the donor, pNPS. BvASST incubated with pNPS (only donor, no acceptor) showed a large proportion of sulfonated ([M+1] + 80) form when compared with BvASST incubated without any substrate (no donor, no acceptor) (Figure 4). Here, we used equimolar concentrations of pNPS and BvASST, aiming to achieve a single-turnover reaction to observe only the unsulfonation of the enzyme in the presence of potential acceptors. If pNPS was in excess, we hypothesized that the enzyme would predominantly remain in its sulfonated form and would interfere with our assay to screen acceptors. However, further analysis revealed that the pNPS concentration was slightly lower than that of BvASST, which likely explains incomplete sulfonation of the enzyme in contrast to full sulfonation observed in Figure 3.

**Fig. 4:**
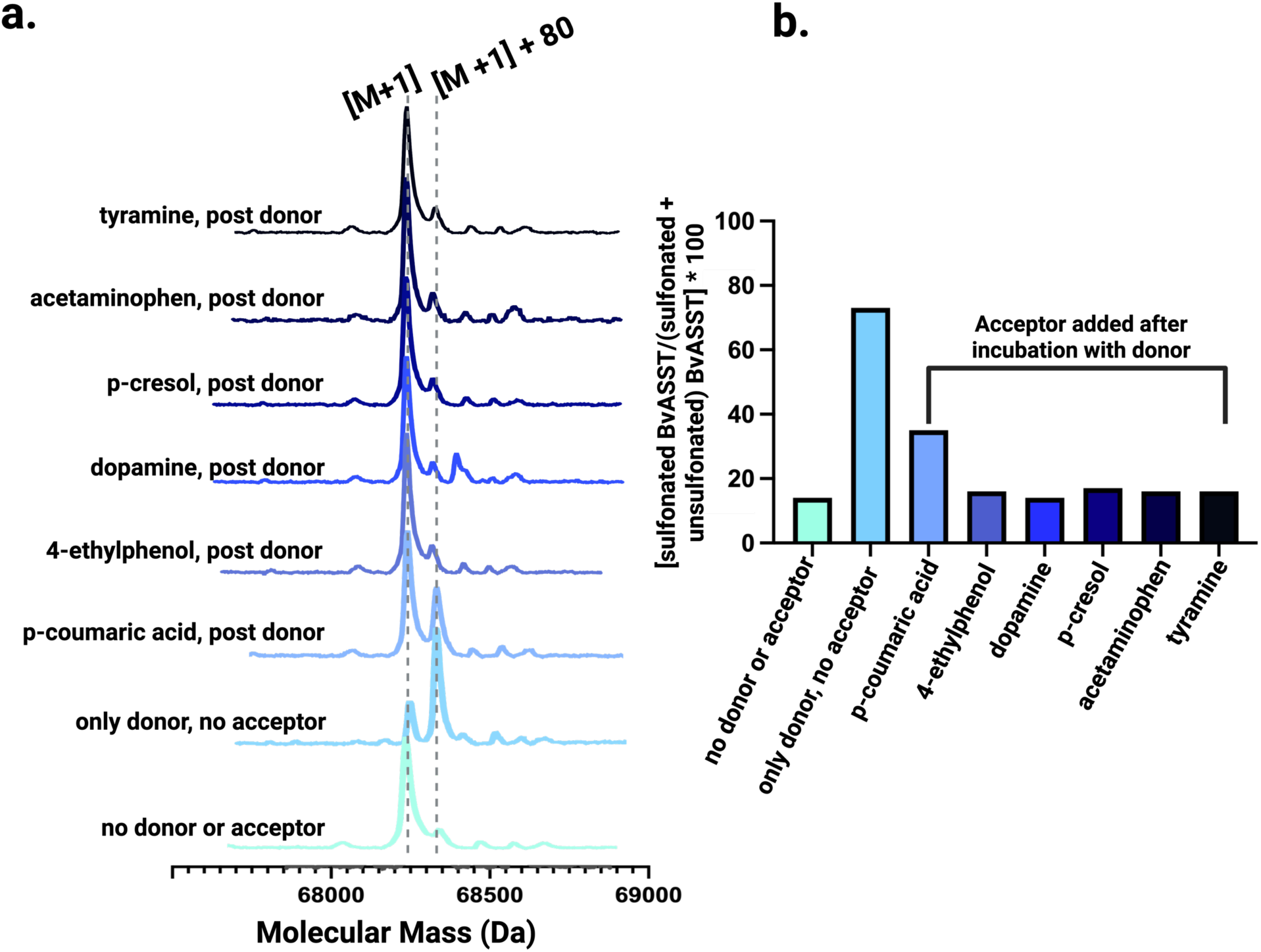
Incubation of sulfonated BvASST with various phenolic molecules reverted it to a largely unsulfonated form, confirming these compounds as compatible sulfo group acceptors. Purified BvASST (no donor or acceptor) incubated with pNPS (20 min) displayed a ([M+1] + 80) Da mass shift (only donor, no acceptor). This sulfonated BvASST, when subsequently incubated with a sulfo acceptor (2 hours), showed a mass shift back to its original mass [M+1]. **a**. Intact mass spectra collected from the incubation of sulfonated BvASST (generated via incubation with pNPS) with each acceptor molecule tested. Controls do not have acceptors added and include: 1. no donor or acceptor which depicts the purified form of BvASST, and 2. incubation with pNPS (sulfo donor) only without any acceptor. **b**. Schematic of qualitative trends for the change in BvASST sulfonation in the presence of compatible phenolic acceptor molecules where colors correspond to spectra shown in panel a, estimated as a percentage of ratio: ([M+1]+ 80)/ ([M+1] + ([M+1]+80]))*100 or [sulfonated BvASST/ (sulfonated + unsulfonated) BvASST]*100.

When sulfonated BvASST was incubated with various sulfo acceptors, we observed a decrease in sulfonated enzyme peak and a corresponding increase in the unsulfonated form (Figure 4a), suggesting possible transfer of the sulfo group from the enzyme to an acceptor molecule. However, this assay does not directly detect the formation of sulfated acceptor products, and the occurrence of sulfonation must be confirmed through additional experimental validation. Figure 4b provides a schematic representation of the qualitative trends observed in the intact mass spectrometry data from Figure 4a. This assay was designed to screen potential sulfo acceptors for BvASST rather than to compare their relative efficiencies. For all tested acceptor molecules, a decrease in enzyme sulfonation was evident (Figure 4a). We observed peak intensity shifts in the presence of tyramine, dopamine, p-cresol, 4-ethylphenol (4-EP), acetaminophen, and p-coumaric acid, indicating that each of these molecules facilitated the conversion of sulfonated BvASST back to its primarily unsulfonated form, making it available for additional catalytic cycles. While we are not quantitatively comparing acceptor efficiencies, p-coumaric acid (pCA) exhibited the smallest decrease in sulfonation, suggesting possible reaction reversibility and equilibrium effects leading to the accumulation of both enzyme forms, though this requires further experimental validation. Collectively, intact mass spectrometry data (Figures 3 and 4) confirmed both half-reactions of BvASST catalyzed sulfo transfer and identified various phenolic compounds as sulfo donors and acceptors for this enzyme.

### BvASST is sulfonated at a histidine residue

To identify the site of sulfonation, BvASST underwent trypsin digestion and the peptide that contained a sulfonated residue was identified by LC-MS/MS (Figure 5). An amino acid residue sulfonated in the reaction catalyzed by BvASST was identified as a histidine residue, specifically H471 (Supplementary Figure 7). This residue is conserved within ASSTs (Supplementary Figure 8) and was identified as the site of sulfonation in an aryl-sulfate sulfotransferase from a uropathogenic strain of an *E. coli*[13].

**Fig. 5:**
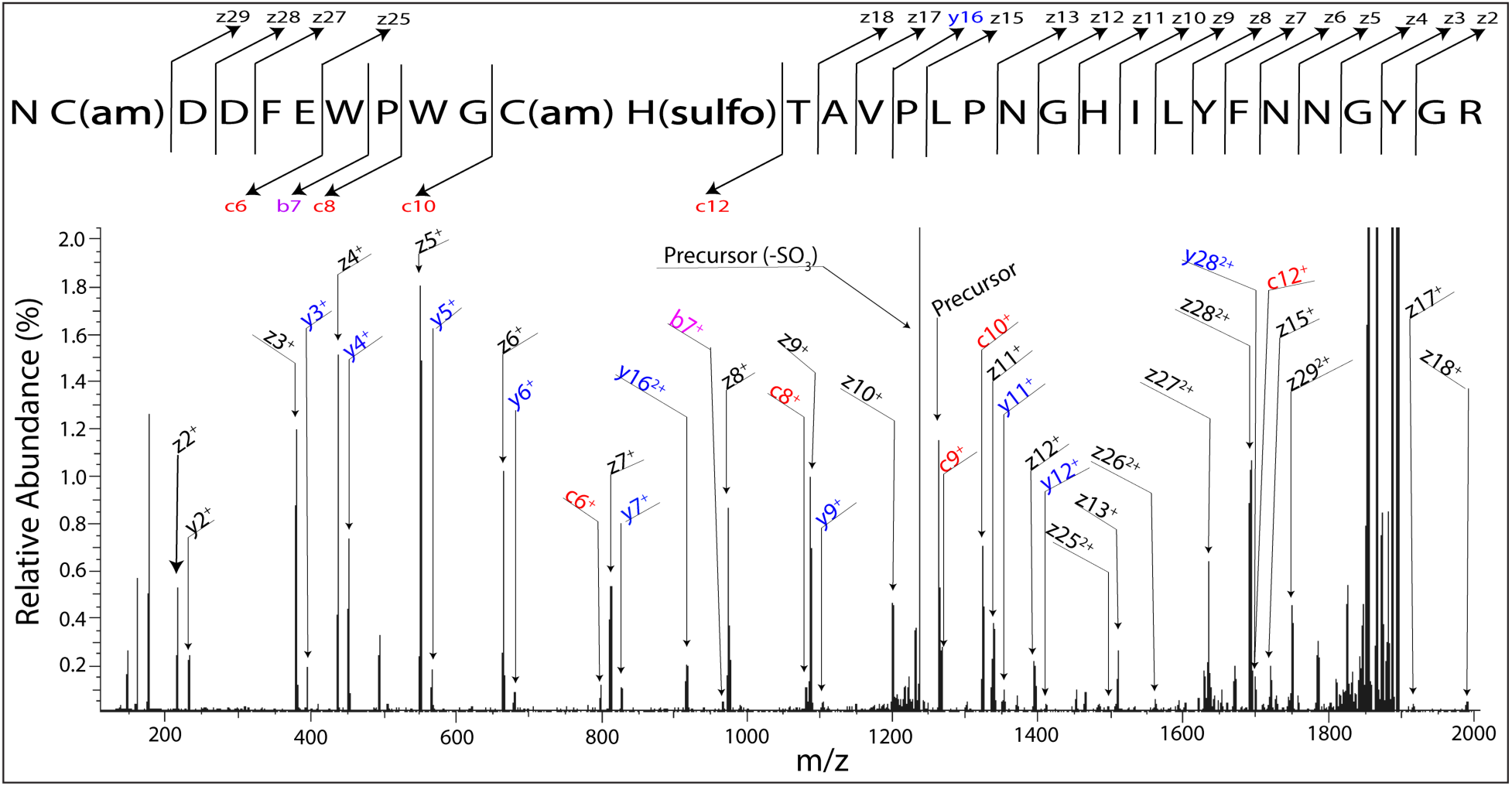
Identification of the site of sulfonation for BvASST as His471. EThcD-MS/MS spectrum of trypsin-generated peptide fragment 460-490. Specific peptide fragments observed in the spectrum are annotated in both the spectrum and the peptide map above it where C(am) indicates alkylated (carb<u>am</u>idomethylated) Cys, which was generated during alkylation of reduced cysteine residues with iodoacetamide. H(sulfo) indicates sulfonated His. Fragments labeled *b* or *c* contain the N-terminus and fragments labeled *y* or *z* contain the C-terminus. From the N-terminal side, the observation of the *c_10+_* ion shown indicates that the first 10 residues are unmodified (other than intentionally alkylated Cys). Observation of the mass-shifted *c_12+_* ion shown indicates that either the C(am) or His(sulfo) residue must contain the sulfonated residue. Since alkylated Cys is not a candidate for sulfonation, the logical conclusion is that the His residue is sulfonated. Similar logic can be applied to the *z*-ion series shown. A spectrum zoomed in on the c12^+^ fragment is provided in the supplemental data (Supplementary Figure 7).

### *In vitro* enzymatic assays with BvASST show sulfo transfer reactions between various donor-acceptor pairs of biologically relevant phenolic compounds

Following the identification of donor and acceptor substrates for BvASST, we tested various donor-acceptor pairs to assess their compatibility and determine if reaction products could be detected. To probe this, we provided BvASST with acetaminophen sulfate, p-coumaric acid sulfate, p-cresol sulfate, dopamine sulfate, PAPS, and p-nitrophenyl sulfate as sulfo donor substrates (Figure 6). We tested each of these sulfo donor substrates with several acceptor substrates that include dopamine, p-cresol, p-coumaric acid (pCA), p-nitrophenol (pNP), and acetaminophen (Figure 6). Interestingly, we detected the production of sulfated phenolic acceptor molecules and desulfated phenolic donor molecules in many reaction pairs (Figure 6a). No products were detected when dopamine sulfate was utilized as a donor with p-cresol and acetaminophen as acceptors. Additionally, we did not observe any sulfated products where PAPS was the sulfo donor, confirming that BvASST is PAPS-independent sulfotransferase. Dopamine, a neurotransmitter, was a universal acceptor for all tested donors and these reactions produced dopamine sulfate (Supplementary Figures 10-15). In contrast, acetaminophen sulfate was found to be the universal donor (Supplementary Figure 14), where sulfonated forms of all phenolic acceptors that include p-coumaric acid, p-cresol, dopamine, and p-nitrophenol were detected along with desulfonated acetaminophen (Figure 6a, Supplementary Figure 14). One such reaction has been depicted in Figure 6b, where acetaminophen sulfate was reacted with p-cresol. In humans, one of the ways to metabolize acetaminophen is via sulfonation and hence acetaminophen sulfate is a xenobiotic metabolite, whereas p-cresol is only produced by gut microbes[35]. Here, we showed that purified BvASST catalyzed a sulfo transfer reaction between acetaminophen sulfate and p-cresol to generate acetaminophen and p-cresol sulfate (Figure 6b). This is an interesting chemistry that might have larger implications. For example, children with ASD have difficulty with the metabolism of the commonly used analgesic and antipyretic agent, acetaminophen[36]. This has partly been attributed to the competition in sulfonation between various acceptors during sulfo transfer reactions catalyzed by human sulfotransferases (hSULTs). It is hypothesized that p-cresol and acetaminophen compete for sulfonation, and that excessive p-cresol sulfate formation interferes with the sulfonation pathway for acetaminophen, one of the principal routes for its metabolism and excretion[37]. These observations highlight the need for a deeper understanding regarding the human gut microbiome’s ability to alter the concentration of these compounds. Our results demonstrate that diverse phenolic molecules and phenolic sulfates which are known to be biologically relevant can be transformed by BvASST *in vitro*.

**Fig. 6:**
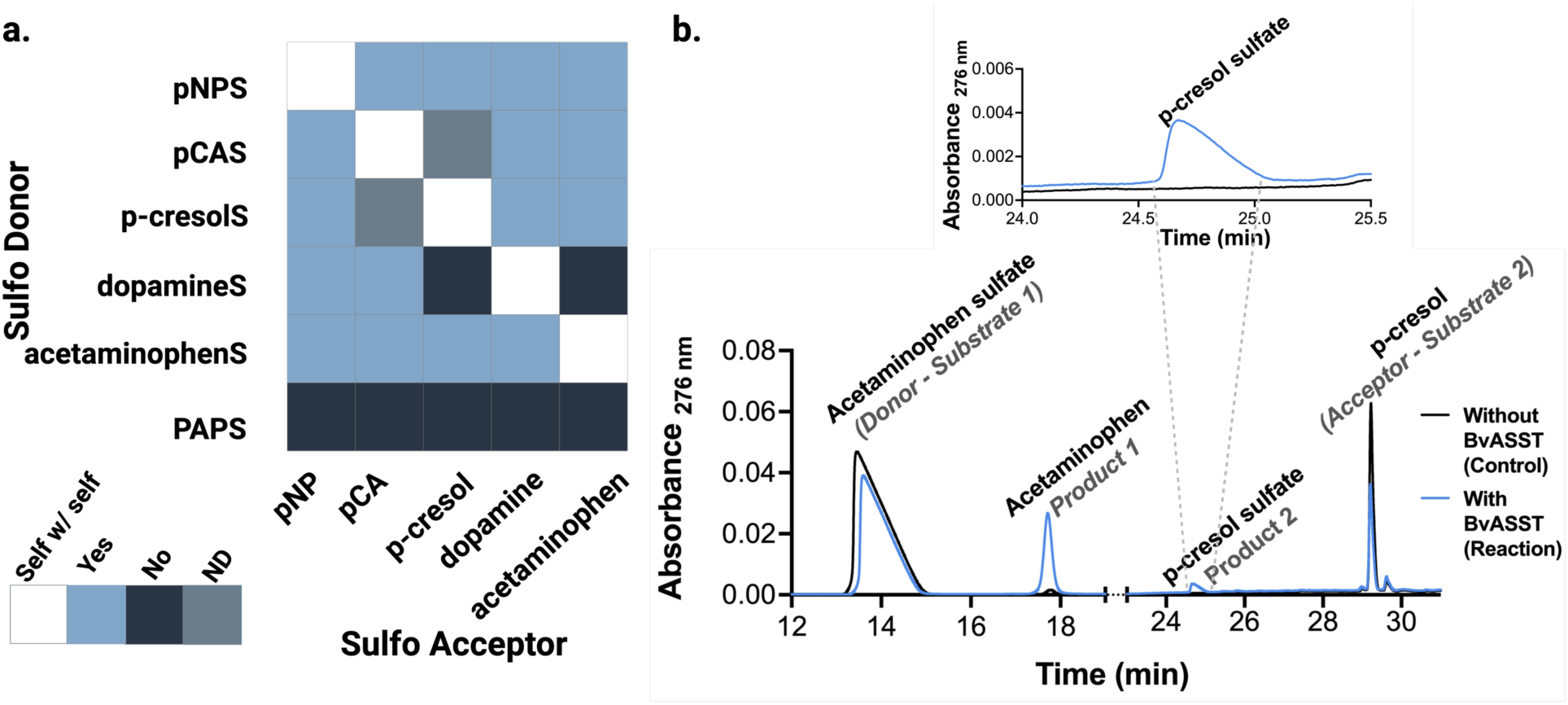
Pairs of substrates (donors and acceptors) react with each other via BvASST catalyzed sulfo transfer reactions. Assays containing various pairs of sulfo group donor and sulfo group acceptor in the presence and absence of BvASST were incubated for 4 hours at 37 °C. After incubations, all assays were analyzed via HPLC to detect products that include sulfated forms of acceptors and desulfated donors which were verified by comparing their retention times to commercial authentic standards. Each substrate pair was replicated three times (n=3) (Supplementary Figures 10-15). **a.** Tested donor and acceptor substrate pair reactions analyzed in terms of generation of products to determine the reaction occurrence. Light blue squares indicate detection of both products in reactions catalyzed by BvASST, dark blue squares indicate no product detection, grey squares indicate inability to separate peaks and therefore the presence of products can’t be determined (ND), and white represents self with self-type reactions (e.g. Acetaminophen sulfate with acetaminophen). All compounds were identified as compared to the retention times of standards for verification. Here, pNPS is p-nitrophenyl sulfate, pCAS is p-coumaric acid sulfate, p-cresolS is p-cresol sulfate, dopamineS is dopamine sulfate, and acetaminophenS is acetaminophen sulfate. **b.** A representative chromatogram depicts a reaction utilizing acetaminophen sulfate as a donor and p-cresol (pC) as an acceptor. In the presence of BvASST, p-cresol sulfate (pCS) and acetaminophen were produced. In control reaction, small peak of acetaminophen was detected due to non-catalytic hydrolysis of small percentage of acetaminophen sulfate (Supplementary Figure 16a), but we do not detect any p-cresol sulfate. A reaction is only positive when both products are detected.

To better understand their interactions with other phenolic molecules, this study also included pNP and pNPS which are synthetic, non-physiological acceptor and donor substrates, respectively. All donor molecules, except PAPS, were able to sulfonate pNP (Supplementary Figures 10-15). Similarly, all phenolic acceptor substrates were successfully sulfonated by the non-physiological donor pNPS (Supplementary Figure 10). Due to their broad reactivity, pNP and pNPS can serve as reliable positive controls in BvASST catalyzed reactions, helping to ensure the experimental setup is functioning optimally.

### Continuous kinetic assays identify potential additional phenolic sulfo group acceptors for BvASST catalyzed reactions

To test if BvASST can utilize additional phenolic molecules as acceptors apart from above tested compounds (Figure 4) *in vitro*, we expanded our library of phenolic acceptors and reacted them with BvASST. Phenolic molecules shown in Figure 7b, including caffeic acid, serotonin, phenol, o-cresol, m-cresol, 2-naphthol, and 4-hydroxyphenylpyruvate (4-HPP) were tested as acceptors for the reaction catalyzed by BvASST. To test these multitude of phenolic acceptors, enzymatic activity assays were conducted with synthetic sulfo donors, p-nitrophenyl sulfate (pNPS) (for colorimetric assays), and 4-methylumbelliferyl sulfate (4-MUS) (for fluorescence-based assays) (Figure 7a, Supplementary Figure 9) which provided continuous, stable, reliable, and high-throughput measurements for all molecules illustrated in Figure 7b[11, 13, 15, 22].

**Fig. 7:**
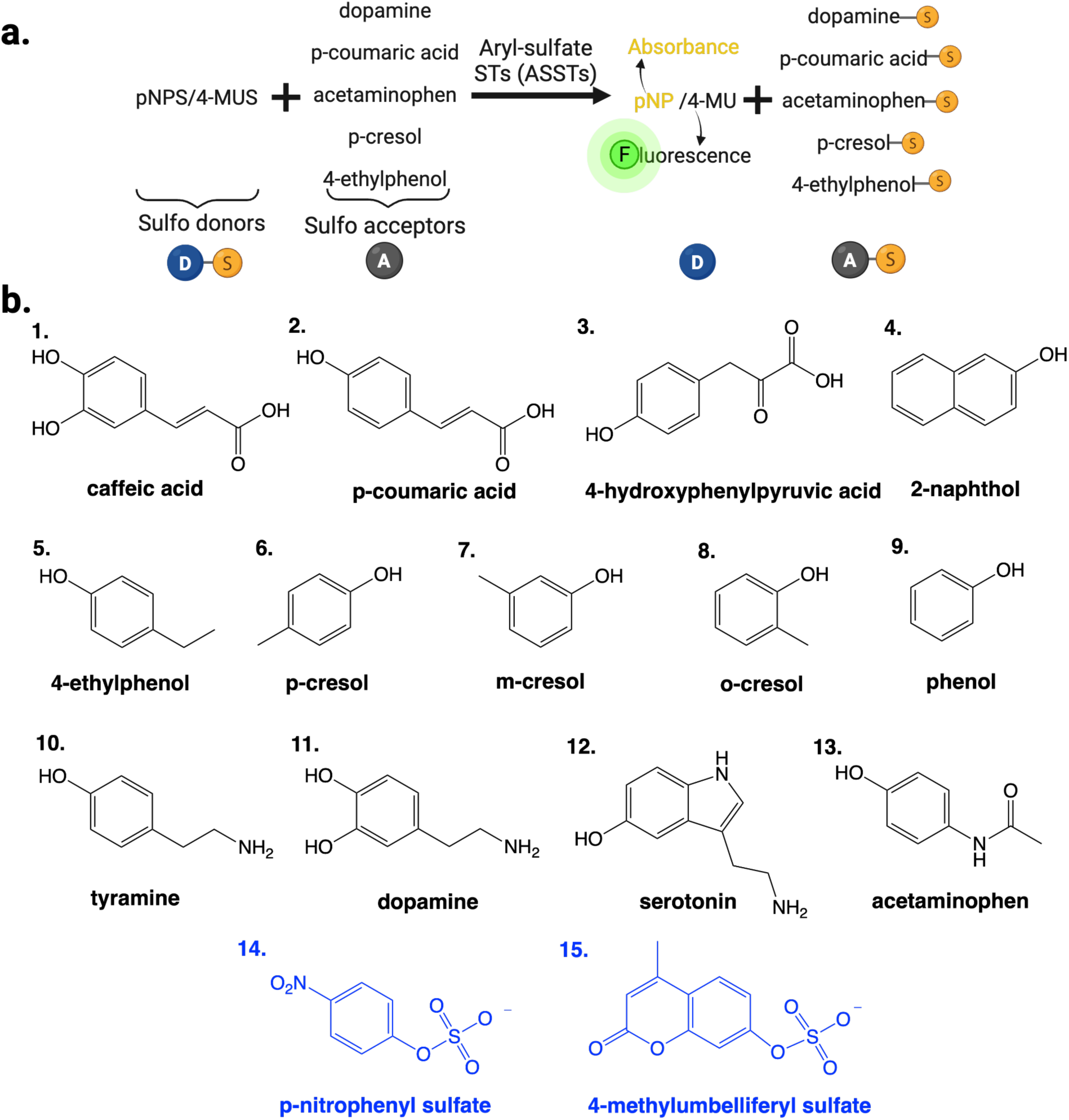
Enzymatic assays to test sulfotransferase activity of BvASST with a variety of phenolic acceptor molecules. **a.** The figure depicts two types of continuous enzymatic assays, a colorimetric assay with pNPS as a sulfo donor and a fluorescence-based assay with 4-MUS as a sulfo donor. **b.** Structures of molecules tested as acceptors with this assay. Blue molecules represent sulfo donors used in kinetic assays depicted in panel a.

When pNPS was used as the sulfo donor, the reaction velocity for most molecules increased linearly with increasing BvASST concentration (Figure 8). This confirmed that the observed product formation was due to enzymatic activity rather than spontaneous or non-catalytic processes and BvASST was able to use these molecules as sulfo group acceptors. For serotonin, little to no activity was observed in the presence of BvASST. Moreover, initial velocities changed significantly depending on the type of the phenolic molecule utilized as an acceptor. The phenolic molecule with the highest velocity was the preferred acceptor for BvASST under our assay conditions. We observed that caffeic acid exhibited the highest reaction rates, followed by p-coumaric acid. Both molecules are structurally similar, differing only by the presence of a hydroxyl group (Figure 7b, structures 1 and 2).

**Fig. 8:**
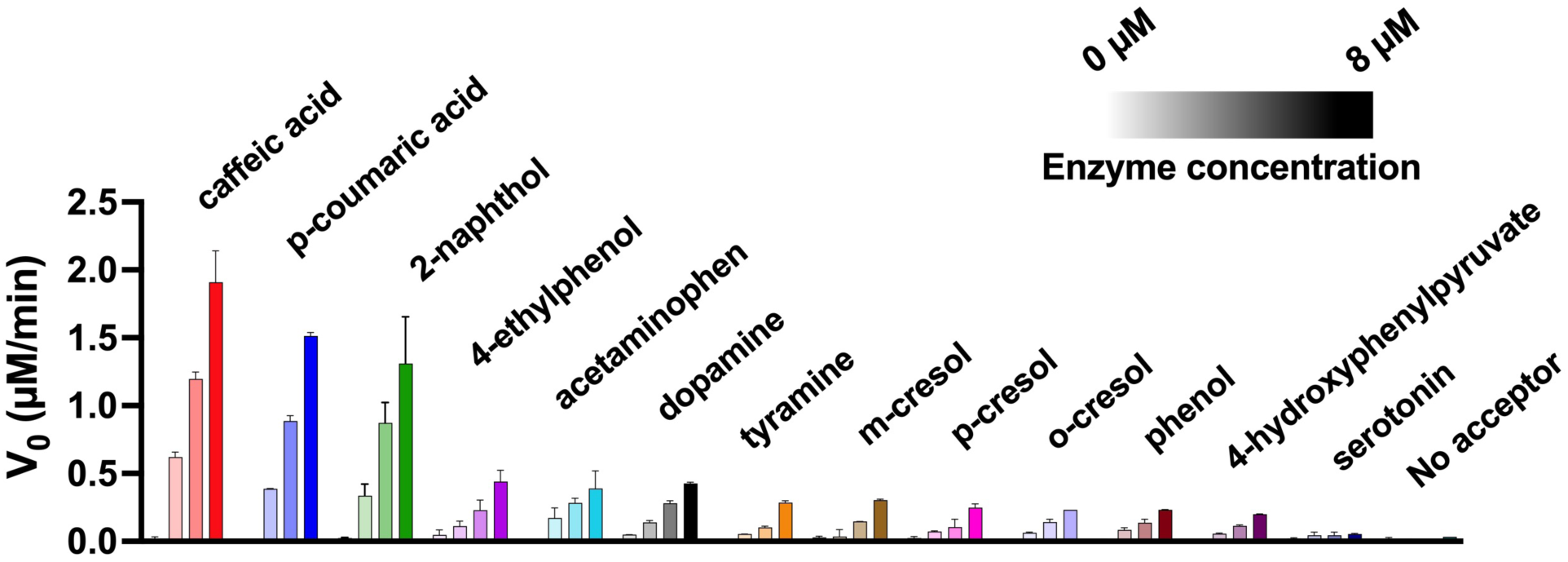
Reaction velocities are proportional to BvASST concentrations for many phenolic acceptors. Initial velocities (V_0_) were measured for BvASST catalyzed reactions with pNPS as a donor in presence of various acceptors, as listed above each dataset. The production of pNP was monitored over time (150 minutes) and from reaction progress curves, initial datapoints were fitted with linear regression to calculate initial velocities. For each acceptor, the shade of the color from light to dark illustrate the increase in BvASST concentration from 0 – 8 µM (0 µM, 2 µM, 4 µM, and 8 µM).

### Ranking of acceptor compatibility remained similar between the two synthetic donors

As described previously, BvASST is capable of utilizing a broad range of acceptor substrates. To evaluate acceptor preference of BvASST, we conducted steady-state kinetic assays (Fig. 7a) in which the sulfo donor was held constant and the acceptor concentration was systematically varied, enabling us to determine kinetic parameters for several sulfo acceptors, including dopamine, acetaminophen, p-cresol (4-methylphenol), 4-ethylphenol, caffeic acid, and p-coumaric acid[11, 14, 15]. The kinetic parameters varied widely for these acceptor molecules. The preference for acceptor was ranked by *k_cat_*/*K_m_*, which describes catalytic efficiency. The relative acceptor ranking for caffeic acid, p-coumaric acid, and dopamine was consistent with either sulfo donor— pNPS or 4-MUS (Tables 1 and 2). For reactions where pNPS was utilized as a donor, acceptors were ranked as: caffeic acid ≥ p-coumaric acid > dopamine. For reactions with 4-MUS as a donor, acceptors were ranked as: caffeic acid ≥ p-coumaric acid > 4-ethylphenol > p-cresol ≥ dopamine. To determine whether acceptor preference is an intrinsic feature of the enzyme, it will be important to test these acceptors with the newly identified phenolic sulfo donors mentioned above whose modulation is known to influence human health. In the case of the BvASST catalyzed reaction between synthetic donors and acetaminophen, the rate of reaction linearly increased with an increase in acetaminophen, with no plateau within the substrate solubility limits (Supplementary File 2). Similarly, a linear trend appeared when pNPS was the donor and either 4-ethylphenol or p-cresol served as acceptors. This absence of saturation could possibly indicate that all tested concentrations were below *K_m_*, or it might hint that the reactions can also proceed appreciably in the reverse direction.

**Table 1:** Steady state kinetic parameters for BvASST with various acceptors using pNPS as a sulfo group donor.

| Acceptor | $K_m$ ($\mu\text{M}$) | $k_{cat}$ ($\text{min}^{-1}$) | $k_{cat}/K_m$ ($\mu\text{M}^{-1} \text{min}^{-1}$) |
| --- | --- | --- | --- |
| caffeic acid<br>n=3 | $22.2 \pm 2.7$ | $0.49 \pm 0.01$ | $0.0220 \pm 0.00240$ |
| p-coumaric acid<br>n=3 | $38.4 \pm 2.3$ | $0.80 \pm 0.06$ | $0.0210 \pm 0.00030$ |
| dopamine<br>n=3 | $2008 \pm 217$ | $0.41 \pm 0.01$ | $0.0002 \pm 0.00002$ |
| 4-ethylphenol<br>n=3 | ND-Unable to saturate | ND- Unable to saturate |  |
| p-cresol<br>n=3 | ND-Unable to saturate | ND- Unable to saturate |  |
| acetaminophen<br>n=3 | ND-Unable to saturate | ND- Unable to saturate |  |

*Kinetic parameters for different sulfo acceptors using pNPS as the sulfo donor. Measured via pNP production by monitoring absorbance at 405 nm ( $\epsilon = 18,000 \text{ M}^{-1} \text{ cm}^{-1}$ ) (Supplementary File 2). Values are expressed as mean ± SD.*

**Table 2:** Steady state kinetic parameters for BvASST with various acceptors using 4-MUS as a sulfo group donor.

| Acceptor | $K_m$ ($\mu\text{M}$) | $k_{\text{cat}}$ ($\text{min}^{-1}$) | $k_{\text{cat}}/K_m$ ($\mu\text{M}^{-1} \text{min}^{-1}$) |
| --- | --- | --- | --- |
| caffeic acid<br>n=3 | 55 ± 21 | 0.045 ± 0.006 | 0.00088 ± 0.000260 |
| p-coumaric acid<br>n=2 | 131 ± 46 | 0.104 ± 0.007 | 0.00085 ± 0.000250 |
| dopamine<br>n=3 | 2292 ± 711 | 0.071 ± 0.016 | 0.00003 ± 0.000002 |
| 4-ethylphenol<br>n=2 | 756 ± 77 | 0.087 ± 0.004 | 0.00012 ± 0.000006 |
| p-cresol<br>n=3 | 2712 ± 169 | 0.101 ± 0.006 | 0.00004 ± 0.000001 |
| acetaminophen<br>n=3 | ND- Unable to saturate | ND- Unable to saturate |  |

*Kinetic parameters for different sulfo acceptors using 4-MUS as the sulfo donor. Measured via 4-MU production by monitoring fluorescence at 460 nm (excitation 360nm) (Supplementary File 2). Concentrations were determined with the help of a standard curve. Values are expressed as mean ± SD.*

## DISCUSSION

Conjugation reactions like sulfonation play a key role in the metabolism and excretion of phenolic compounds. Many biologically important metabolites fall into this category, including phenolic sulfates. Altered levels of phenolic compounds have been linked to various disorders[38–40]. For instance, studies have reported gut microbial dysbiosis and changes in both sulfated and non-sulfated phenolic compounds in the fecal samples of children with autism spectrum disorder (ASD)[41, 42]. Phenolic molecules like p-cresol and phenol are known to be differentially abundant between children with ASD and typically developing (TD) children[19, 41–43]. Moreover, sulfated derivatives of phenolic compounds like p-cresol sulfate (pCS) and 4-ethylphenyl sulfate (4-EPS), have been shown to exert toxic or adverse effects on the host[44–46]. For example, 4-ethylphenyl sulfate (4-EPS) has been linked to anxiety-like behaviors in mice[44]. Dopamine, a neurotransmitter produced in large amounts in the gastrointestinal tract[47], has also been implicated in the pathophysiology of Alzheimer’s disease[48], Parkinson’s disease[49], and obesity[50]. Therefore, understanding the factors that regulate sulfonation of these phenolic compounds is crucial.

To investigate the potential role of the human gut microbiome in the sulfonation of small phenolic compounds, we performed a bioinformatics analysis to identify putative genes annotated to encode aryl-sulfate sulfotransferase (ASST) enzymes which catalyze sulfo group transfer reactions involving phenolic substrates, within prevalent gut microbial species. To probe catalytic function and to identify substrates, we selected a representative aryl-sulfate sulfotransferase (BvASST) from the abundant gut microbe *Bacteroides vulgatus*, according to the criteria outlined earlier. Interestingly, many *Bacteroides* species contain an additional sulfotransferase gene encoding a SULT homolog, which is believed to function as a PAPS-dependent sulfotransferase[9, 10]. Unlike most *Bacteroides* species, *B. vulgatus* is unique in harboring two distinct sulfotransferase genes, one encoding a potential PAPS-dependent sulfotransferase as previously reported in *B. thetaiotaomicron* [9, 10], and the other annotated as an aryl-sulfate sulfotransferase (BvASST). The reason for the coexistence of these two sulfotransferases in *B. vulgatus* and their respective contributions to microbial and host physiology remains unclear. Interestingly, we found that when expressed in *E. coli*, BvASST functions as a lipoprotein which is an unusual characteristic, as none of the previously studied ASSTs have been identified as lipoproteins[9, 11–14, 16, 17, 19, 32, 43].

With this study, we demonstrated that BvASST can utilize a wide range of structurally diverse phenolic compounds as sulfo group acceptors. These include phenolic acids (e.g., caffeic acid, p-coumaric acid, 4-hydroxyphenylpyruvate); phenolic amines (e.g., dopamine, tyramine); phenolic amides (e.g., acetaminophen); simple phenolics (e.g., phenol, p/o/m-cresols, 4-ethylphenol); and a bicyclic phenol (2-naphthol). Interestingly, these molecules vary not only in their chemical classifications but also in their sources of origin. For example, we observed catalysis with dietary compounds like caffeic acid and p-coumaric acid; microbially produced compounds such as p-cresol, p-coumaric acid, and 4-ethylphenol; synthetic xenobiotics like acetaminophen; host or microbiota-derived molecules such as tyramine, dopamine, and phenol[43]; host metabolic intermediates like 4-hydroxyphenylpyruvate (4-HPP); and naturally occurring compounds like 2-naphthol, a known precursor to several human-associated xenobiotics[44]. These findings highlight the catalytic versatility of BvASST, which is capable of recognizing and modifying a broad spectrum of phenolic substrates *in vitro* with diverse chemical structures and biological origins.

Unlike PAPS-dependent sulfotransferases such as cytosolic human sulfotransferases (hSULTs) and microbial SULTs which utilize 3’-phosphoadenosine-5’-phosphosulfate (PAPS) as a sulfo group donor, the identities of sulfo donors for aryl-sulfate sulfotransferases (ASSTs) have remained largely unknown. Using intact mass spectrometry and HPLC, we identified several sulfo group donors for BvASST *in vitro*.

These included biologically relevant sulfated metabolites such as acetaminophen sulfate, dopamine sulfate, p-coumaric acid sulfate, and p-cresol sulfate, compounds whose levels are known to influence host health. If such sulfo transfer reactions occur in the gut environment, it raises the possibility that gut microbial ASSTs may contribute to or modulate host metabolite pools. For instance, the sulfonation of acetaminophen is generally attributed to human sulfotransferases for its detoxification and excretion[43]. However, our results show that BvASST can catalyze a similar reaction *in vitro*, suggesting that gut microbes may either complement or interfere with this host detoxification pathway. If this activity occurs *in vivo*, it could potentially alter the pharmacokinetics of acetaminophen by affecting its clearance rate and influencing optimal dosing, as a result of gut microbial enzyme activity. In such cases, microbial contributions to acetaminophen metabolism could have important implications for drug efficacy and toxicity[44]. However, further *in vivo* studies are needed to experimentally validate this hypothesis.

Consistent with other characterized ASSTs, BvASST was capable of utilizing synthetic sulfo donors that are not naturally found in biological systems or environmental contexts. While these donors are unlikely to be encountered by the enzyme *in vivo*, they serve as useful tools for assessing enzymatic activity due to the colorimetric or fluorometric properties of their non-sulfated counterparts. The ability of BvASST to act on such non-natural substrates likely reflects coincidental rather than evolutionarily selected activity. This observation further suggests a degree of active site plasticity, enabling BvASST to accommodate and transform phenolic molecules that are potentially non-native to the organism.

In conclusion, we identified a microbial sulfotransferase from a human commensal gut microbe that functions as a versatile catalyst. BvASST accepts a broad range of phenolic substrates and utilizes previously unrecognized sulfo donors to modulate the levels of phenolic metabolites *in vitro* that have known relevance to the gut-brain axis and overall human health. Future studies should aim to elucidate the physiological significance of these BvASST mediated transformations *in vivo*.

## Author Contribution

DS, RKB, and RC conceptualized the experiments. RC performed majority of the experiments. SC and CRB performed the mass spectrometry experiments, that included method development and data analysis. MM developed HPLC method and RC and MM collected the HPLC data. AB performed the inorganic sulfate analysis. KN performed shotgun metagenomics data analysis. RC and DS wrote the manuscript. RKB, SC, CRB, MM, AB, KN reviewed the manuscript and provided edits.

